# Expression patterns of Arabidopsis *bZIP19* and *bZIP23* elucidate their individual contribution to the regulation of the Zn deficiency response

**DOI:** 10.1101/2025.07.03.662962

**Authors:** Ana Campilho, Grmay Hailu Lilay, Margarida Benvindo, Pedro Humberto Castro, Ana G. L. Assunção

## Abstract

Plants control the availability of nutrients through tightly regulated homeostasis networks. In *Arabidopsis thaliana*, the transcription factors bZIP19 and bZIP23 regulate the homeostasis of the micronutrient Zn, acting as the key regulators of the Zn deficiency response and as sensors of cellular Zn status. Although there is a degree of functional redundancy between bZIP19 and bZIP23, their individual contributions in this regulatory process are not yet understood. To gain insight on the individual contributions of bZIP19 and bZIP23, we analysed the phenotypes of *bzip19, bzip23* and *bzip19bzip23* single and double mutants in response to Zn deficiency, and histochemical GUS staining in *pbZIP19::GUS* and *pbZIP23::GUS* lines. Analysis of mutant phenotypes indicated unequal redundancy between bZIP19 and bZIP23. Expression of GUS in *pbZIP19::GUS* was observed in differentiated root tissues including root hairs, in the vascular tissue of hypocotyl, cotyledons and petals, in leaf trichomes and stamen filament. In *pbZIP23::GUS*, the expression was predominant in root and shoot apical meristems, being also observed in leaves, root vasculature, stamen filament and anthers. In summary, *bZIP19* and *bZIP23* have different and mostly non-overlapping tissue-specific expression patterns. This may explain the unequal redundancy between bZIP19 and bZIP23 and their evolutionary retention as paralogs.

## INTRODUCTION

Micronutrients are required by plants, at adequate concentrations, fo**r** optimal growth. Therefore, in order to avoid deficiency or excess, plants rely on a tightly regulated micronutrient homeostasis network. The prevalence of micronutrient deficiencies in cultivated soils and plants are a global problem that negatively affect crop production and plant nutritional value, as well as human health and well-being (Assunção *et al*. 2022).

In the model plant *Arabidopsis thaliana* (Arabidopsis), the zinc (Zn) homeostasis network is regulated by the transcription factors (TFs) bZIP19 and bZIP23 (Assunção *et al*. 2010). These are the key regulators of the response to Zn deficiency, and they also act as sensors of cellular Zn status (Lilay *et al*. 2021). bZIP19 and bZIP23 are members of the basic region leucine-zipper (bZIP) family of eukaryotic transcription factors (Dröge-Laser *et al*. 2018). They belong to the Arabidopsis F group bZIPs (F-bZIP), characterized by a conserved N-terminus cysteine- and histidine-rich motif identified as Zn Sensor Motif (ZSM). In response to Zn deficiency, bZIP19/23 transcriptionally activate the expression of target genes involved in Zn homeostasis. These include Zn transporter genes from the ZIP (ZRT/IRT-like Protein) family of divalent metal transporters (*AtZIP1/3/4/5/9/10/12,IRT3*), which mediate the influx of Zn into the cytosol, and *NAS* genes (*AtNAS2/4*), which encode the enzyme producing nicotianamine, a ligand that forms complexes with Zn to mobilize Zn in the plant (Clemens 2019; Assunção 2022), thus, the Zn-deficiency induced target genes encode Zn transporters and Zn ligands. When there is sufficient Zn supply, the bZIP19/23 TFs act as sensors of the cellular Zn status, through binding of Zn ions to their ZSM, which halts transcriptional activation of target gene expression and the response to Zn deficiency (Lilay *et al*. 2021).

In addition to bZIP19 and bZIP23, which share 69% amino acid sequence identity, there is a third F-bZIP member, bZIP24, sharing 28% and 32% with bZIP19 and bZIP23, respectively. bZIP24, however, does not seem to play a role in the Zn deficiency response (Lilay *et al*. 2019) and instead is reported to regulate salt stress tolerance (Yang *et al*. 2009). The Arabidopsis *bzip19 bzip23* double mutant (*bzip19/23*) displays a Zn deficiency hypersensitive phenotype when growing at Zn deficient conditions. This hypersensitive phenotype is functionally complemented by constitutive expression of either *bZIP19* or *bZIP23* (Assunção *et al*. 2010), including restoring the target genes expression to the Zn-responsive wild-type pattern (Lilay *et al*. 2019), but not by expression of *bZIP24* (Lilay *et al*. 2019). This indicates functional redundancy between bZIP19 and bZIP23 transcription factors. However, the analysis of single mutants grown under Zn deficiency showed a Zn deficiency phenotype in *bzip19*, less severe than in *bzip19/23*, and no visible phenotype in *bzip23*, which indicates that bZIP19 and bZIP23 are not fully redundant (Assunção *et al*. 2010).

The individual contributions of bZIP19 and bZIP23 in the regulation of the Zn deficiency response are not yet elucidated, and it is important to advance the mechanistic understanding of the Zn sensing and deficiency response in Arabidopsis. To obtain insight on their individual contributions to the Zn deficiency response, we studied *bZIP19* and *bZIP23* gene expression patterns by performing a comparative analysis of tissue-specific expression patterns using the *pbZIP19::GUS* and *pbZIP23::GUS* Arabidopsis lines.

## MATERIAL AND METHODS

### Plant material

The Arabidopsis lines used in this study were wild type (Col-0) and the single and double mutant lines *bzip19, bzip23* and *bzip19 bzip23* (*bzip19/23*). The single mutants correspond to the T-DNA insertion mutants *bzip19-1* and *bzip23-1* of the *bZIP19* (At4g35040) and *bZIP23* (At2g16770) transcription factor genes, respectively, and the double mutant *bzip19/23* was selected by crossing between *bzip19-1* and *bzip23-1*, as previously described (Assunção *et al*. 2010). For the histochemical assay, Arabidopsis transgenic lines expressing promoter-β-glucuronidase (GUS) fusion were generated. The promoter regions (the whole intergenic region upstream the start codon) of *bZIP19* and *bZIP23* were amplified from Arabidopsis genomic DNA and cloned into the NotI/AscI restriction enzyme sites of the pENTR™/D-TOPO vector, followed by in vitro site-directed recombination into pBGWFS7 Gateway vector (Karimi *et al*. 2002) carrying a C-terminal GFP fluorophore and GUS reporter gene, using LR Clonase™ II Enzyme Mix (Invitrogen), as described by Lilay et al. (2019). Transformation of *Agrobacterium tumefaciens* and Arabidopsis were performed as described previously (Lilay *et al*. 2019). Transgenic plants were selected by Basta resistance, and seeds (T2 generation) of several independently transformed lines per construct were obtained. These lines were referred to as *pbZIP19::GUS* and *pbZIP23::GUS* lines.

### Growth conditions

Arabidopsis wild-type (Col-0), *bzip19, bzip23* and *bzip19/23* lines were grown in hydroponics in a 10 L container with a modified half-strength Hoagland nutrient solution containing: 2 mM Ca(NO_3_)_2_, 1 mM MgSO_4_.7H_2_O, 3 mM KNO_3_, 1 mM KH_2_PO_4_, 25 µM Fe-Na-EDTA, 25 µM H_3_BO_3_, 3 µM MnSO_4_.H_2_O, 0.1 µM CuSO_4_.5H_2_O, 0.5 µM (NH_4_)_6_Mo_7_O_24_, 50 µM KCl, with 1 mM MES at pH 5.7, as previously described in Lilay et al. (2019). Seeds were germinated on 1.5 mL plastic microtubes, filled with 0.8% (w/v) phyto-agar solution and placed in a support on the hydroponics container (Lilay *et al*. 2019). Plants grew for 6 weeks with a control solution containing 2 μM ZnSO_4_ (control) or with a Zn-deficient solution with no added Zn (-Zn), in a growth chamber with 8/16 h light/dark cycle, 125 μmol m^−2^ s^−1^ white light, 70% relative humidity and 22/19°C light/dark temperature. The nutrient solution was replaced once a week in the first week and twice a week thereafter. For the plate assays, seeds were stratified for 2 days at 4ºC, in the dark, and sterilized with ethanol 70% (v/v), followed by sodium hypochlorite 20% (v/v) and several washes with ultrapure sterile water. Seeds were germinated on petri dishes with ½ MS medium (control) or ½ MS medium with no added Zn (Zn-). Plates were placed vertically in a growth chamber for 7, 12 or 14 days with the conditions described above.

### Tissue element analysis (ICP-OES)

Four-week-old plants, hydroponically grown at control or -Zn nutrient solution, were harvested. Rosette leaves and roots from 5 plants per line (Col-0, *bzip19, bzip23* and *bzip19/23*) and per treatment (control, -Zn) were harvested separately. Roots were desorbed with ice-cold 1 mM CaCl_2_ solution for 5 min followed by 3 times washing with ultrapure water and blotted dry. Leaves and roots were dried for 3 days at 60 °C and their dry weight measured. Samples were digested in a pressurized microwave oven (Ultrawave, Mile-stone Inc.) using ultra-pure acids (70% HNO3, 15% H2O2) at 240°C and 8000 kPa for 15 min. Prior to elemental analysis, samples were diluted to a concentration of 3% HNO3. The multi-elemental analysis was performed with a inductively coupled plasma optical emission spectrometer (5100 ICP-OES, Agilent Technologies). The ICP-OES was equipped with a SeaSpray nebulizer and a double-pass Scott-type spray chamber. Automatic sample introduction was performed from a ASX-520, CETAC auto-sampler from 15 ml falcon tubes. Certified reference material (NIST1515, apple leaf, National Institute of Standards and Technology) was included to evaluate the data quality and drift correction was performed on the basis of drift samples for every 20 samples. Data were processed using the Agilent ICP Masshunter Software (version 4.3) and Agilent ICP Expert Software (version 7.3).

### Histochemical staining for β-glucuronidase (GUS) assay and imaging

Seven-12- and 14-day-old seedlings of Arabidopsis *pbZIP19::GUS* and *pbZIP23::GUS* lines were grown on plates with ½MS medium. Seedlings (*ca*. 4) from four to six independently transformed T2 generation lines from *pbZIP19::GUS* and *pbZIP23::GUS* were analysed. Flowers and siliques from soil-grown *pbZIP19::GUS* and *pbZIP23::GUS* plants, from the above mentioned independently transformed lines, were analysed. Seedlings, flowers and siliques were immersed in GUS staining solution containing 50 mM phosphate buffer, 10 mM Na_2_-EDTA, 20% (v/v) methanol, 0.1% (v/v) Triton X-100, 1.4 mM K3[Fe(CN)6], 1.4 mM K4[Fe(CN)6].3H2O with 1.9 mM X-Gluc, and were incubated overnight at 37°C in the dark (Jefferson *et al*. 1987). After incubation, the pigments were removed by successive incubations in 50%, 70% and 96% (v/v) ethanol, and materials stored in 70% (v/v) glycerol. DIC optics and/or bright field images of GUS-stained seedlings, flowers and siliques were recorded with a Leica Digital Microscope DM6000 equipped with a digital camera Leica MC170 HD, and with an Olympus SZX12 Stereo Microscope equipped with an Olympus DP21 Microscope Camera with U-CMAD3 adaptor.

### Statistical analysis

To compare lines or treatments we used one-way analysis of variance followed by Tukey post hoc tests calculated with IBM SPSS Statistics v 30.0.0.

## RESULTS

### bZIP19 and bZIP23 show unequal redundancy in their response to Zn deficiency

To investigate the individual contribution of bZIP19 and bZIP23 transcription factors in the regulation of the Zn deficiency response, the single and double *bzip19* and *bzip23* T-DNA insertion knock-out mutants (Assunção et al., 2010) were grown for 6 weeks in hydroponics with control (with 2 µM Zn) or Zn deficient (no added Zn) nutrient solution (**Fig. 1; Fig. S1**). The plants grown with control conditions showed no visible differences between genotypes, contrary to the plants grown under Zn deficiency (Zn-) that showed pronounced differences: the growth of *bzip19/23* double mutant was strongly impaired, the *bzip19* single mutant had also strong growth reduction but less severe than *bzip19/23* and, the *bzip23* single mutant had no visible difference compared to the wild-type (**Fig. 1A**). These observed phenotypes were in agreement with previous reports (Assunção *et al*. 2010; Lilay *et al*. 2019). The shoot dry weight showed generally a similar pattern to that of the rosette images, having significantly reduced values for *bzip19* and more severely for *bzip19/23* at Zn deficiency compared with the control conditions, and with the other genotypes (**Fig. 1B**). A similar pattern is also observed in the roots, showing overall an agreement between the dry weight data and the rosette images in the mutant phenotype analysis. Due to the very small size of *bzip19/23* roots grown at Zn deficiency, there is no data for this combination of genotype and treatment (“n.d.” in **Fig. 1B**). The analysis also showed increased shoot and root dry weight at Zn deficiency compared with control supply in wild-type and *bzip23* (**Fig. 1B**), as has been previously observed for the wild-type in a similar hydroponic experiment (Lilay *et al*. 2019).

**Figure 1.**
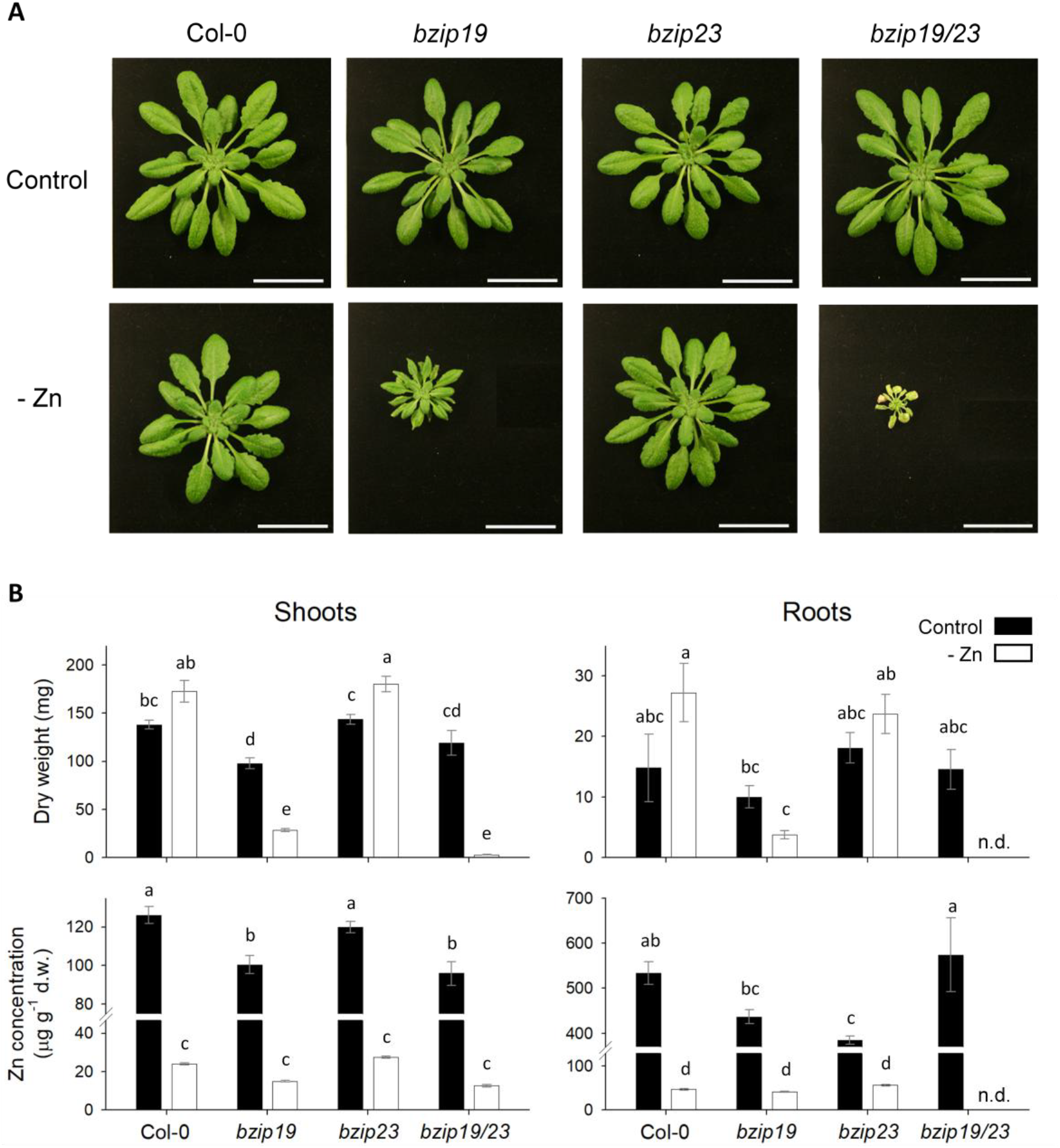
Phenotype of *bzip19, bzip23, bzip19/23* and Col-0 Arabidopsis plants grown in hydroponics with control or Zn deficient (Zn-) nutrient solution. (**A**) Representative image of rosettes from 6-week-old plants. Scale bar = 3 cm. (**B**) Shoot (rosette) and root dry weight (d.w.) (mg), mean ± s.e., n= 3 to 5, and Zn concentration (µg g^-1^ d.w.) measured by ICP-OES, mean ± s.e., n= 3 to 5. There is no data (n.d.) for *bzip19/23* roots under Zn deficiency. Different letters indicate significant differences (P < 0.05), determined using one-way analysis of variance followed by Tukey post hoc tests.

As expected, the Zn concentration in shoots and roots at Zn deficiency was overall strongly reduced compared with the control treatment (**Fig. 1B**). The values of Zn concentration in shoots and roots of the *bzip19/23* and wild-type lines that we observed were comparable to those reported in a previous similar hydroponic experiment (Lilay *et al*. 2019). Although the analysis of *bzip23* did not show a visible mutant phenotype in dry weight or Zn concentration (**Fig. 1**), the ratio of shoot/root Zn concentrations at control supply, but not at Zn deficiency, was slightly higher in *bzip23* than the wild-type suggesting a difference in Zn allocation pattern in this mutant (**Fig. S2**). At Zn deficiency, *bzip19* and *bzip19/23* showed lower Zn concentration in shoots, with average values of 14.96±0.36 and 12.66±0.66 µg g^-1^ dw (mean±s.e.), compared to *bzip23* and wild-type, with 27.42±0.53 and 23.98±0.57 µg g^-1^ dw. Interestingly, these concentration values and the corresponding phenotypes (**Fig. 1A**) are in agreement with the reported threshold of 15–20 µg Zn g^−1^ dw in leaves, below which there is Zn deficiency (Marschner 2012), showing an alignment between the concentration threshold and the onset of Zn deficiency symptoms. The shoot/root Zn concentration ratio in *bzip19* at Zn deficiency, not measured in *bzip19/23*, was smaller than *bzip23* and wild-type suggesting a reduced translocation to shoot (**Fig. S2**).

The comparative analysis of phenotypes in *bzip19, bzip23* and *bzip19/23* showed that, in *bzip23* mutant plants, the activity of the remaining bZIP19 transcription factor seems sufficient to compensate for the loss-of-function of the *bZIP23* gene, whereas in *bzip19* mutant plants the activity of the remaining bZIP23 transcription factor does not compensate for the loss-of-function of the *bZIP19* gene. In addition, the *bzip19/23* double mutant plants displayed enhanced severity of the phenotypes, which indicates that the two transcription factors bZIP19 and bZIP23 are necessary for optimal regulation of the Zn deficiency response, and that they have unequal functional redundancy regarding this regulatory process.

### Analysis of *pbZIP19::GUS* and *pbZIP23::GUS* seedlings shows different *bZIP19* and *bZIP23* tissue-specific expression

To investigate the expression pattern of *bZIP19* and *bZIP23* genes in Arabidopsis, the promoter-GUS fusion lines *pbZIP19::GUS* and *pbZIP23::GUS* were analysed. The histochemical GUS staining of these lines was reported earlier (Lilay *et al*. 2019), showing that *bZIP19* was mostly expressed in the roots whereas *bZIP23* was expressed mostly in the shoot, irrespectively of growth Zn supply. This Zn-independent GUS expression was aligned with *bZIP19* and *bZIP23* transcript levels, which showed no major differences between control or Zn deficient growth conditions (Lilay *et al*. 2019). Here, we analysed expression patterns of *pbZIP19::GUS* and *pbZIP23::GUS* lines in detail. Seven and 12-day-old seedlings were grown on ½ MS (control) medium prior to histochemical analysis (**Fig. 2-4**). In addition, seven and 14-day-old seedlings were grown on Zn deficient ½ MS (Zn-) medium prior to analysis (**Fig. S3**).

**Figure 2.**
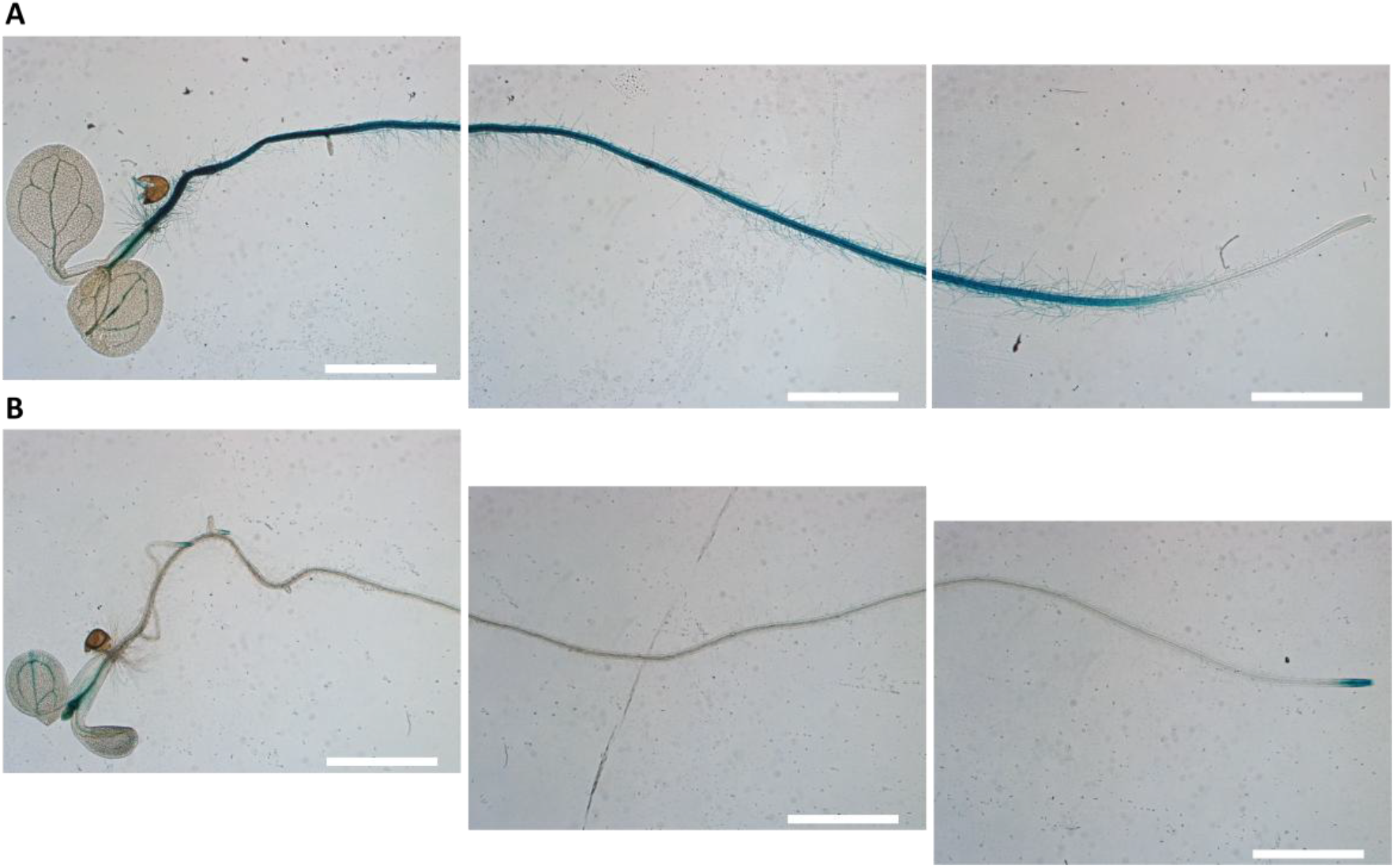
Overview of Arabidopsis seedlings after histochemical GUS staining. Seven-day-old seedling grown on MS (control) from *pbZIP19::GUS* (**A**) and *pbZIP23::GUS* (**B**). The overview is composed with three combined images of the same seedling. Scale bar = 2 mm.

An overview of the histochemical GUS staining of seven-day-old seedlings (**Fig. 2**) showed *bZIP19* expressed mainly in the root, and *bZIP23* expressed in the cotyledons, shoot apical meristem and root tip. More detailed analysis of *pbZIP19::GUS* seedlings (**Fig. 3**) showed no GUS expression in the meristematic and elongation regions of the root, nor in the lateral root primordia (**Fig. 3A-D**). GUS expression only becomes visible in the root differentiation zone where root hairs are present, with the latter being also clearly stained. This was observed in both seven- and 12-day-old seedlings (**Fig. 3B, I, J**). In the root differentiation zone, *pbZIP19::GUS* expression seems present in all root layers (epidermis, cortex and vascular cylinder) (**Fig. 3D,J**), but root cross-section analysis will be needed to confirm this. At the hypocotyl, GUS expression seems restricted to the vascular cylinder (**Fig. 3E,F**) and, in the cotyledons, it appears only in the vascular tissue (**Fig. 3G**,**L**). Conversely, the shoot apical meristem and leaf primordia showed no GUS expression (**Fig. 3E,H**). Interestingly, the trichomes in the leaves have clear GUS staining (**Fig. 3M**). This analysis showed that *bZIP19* is strongly expressed in differentiated root tissues, including root hairs, in vascular tissue of hypocotyl and cotyledons, and in leaf trichomes.

**Figure 3.**
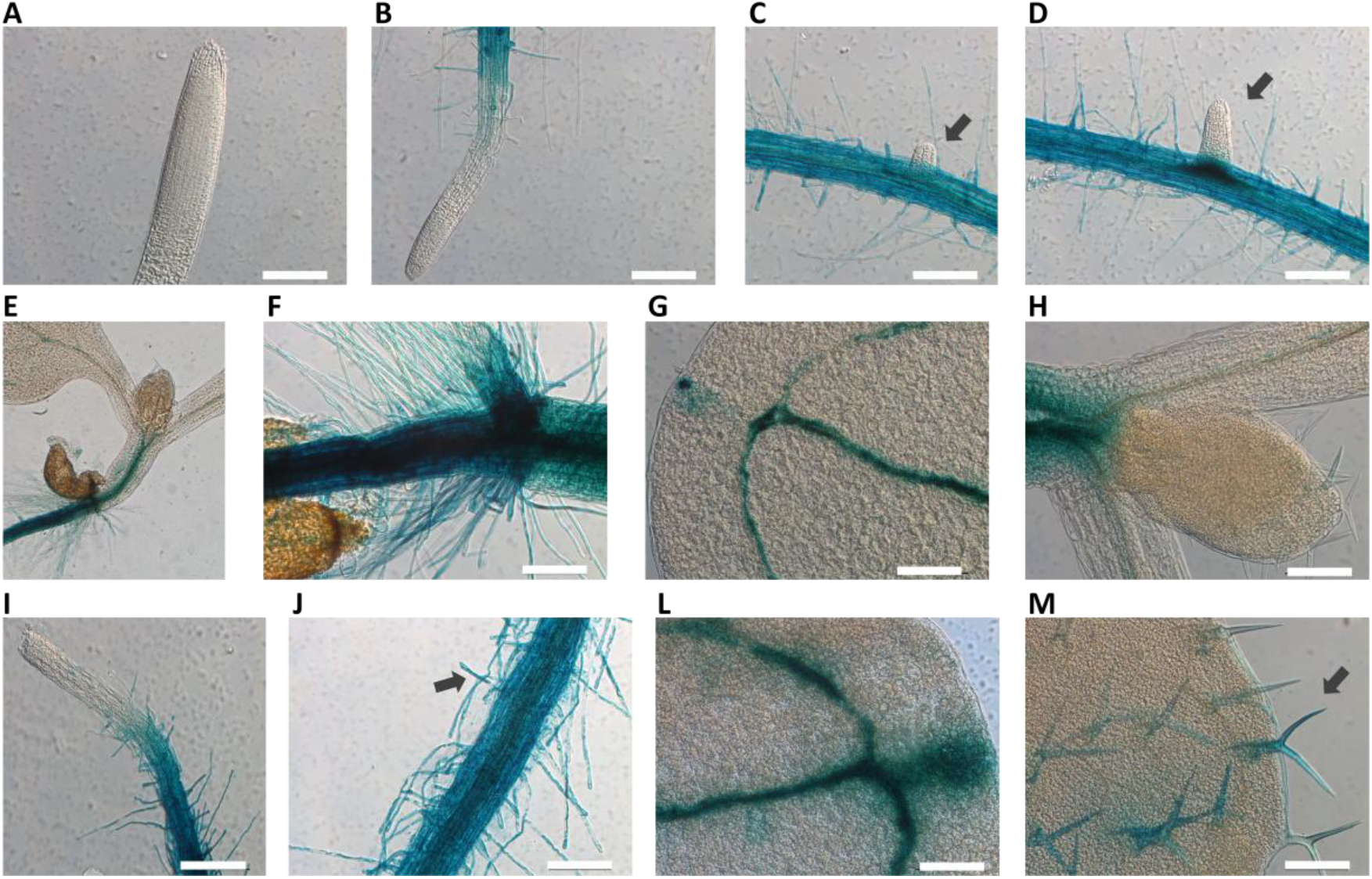
Analysis of *pbZIP19::GUS* seedlings after histochemical GUS staining. Seven-day-old seedlings grown on MS (control) showing the root apical meristem (**A**); root meristem and elongation zone (**B**); lateral root primordia (**C**,**D**); root-hypocotyl junction (**E**,**F**); cotyledon (**G**); shoot apical meristem and leaf primordia (**H**); 12-day-old seedlings showing root apical meristem (**I**); root differentiation zone with root hairs (**J**); cotyledon (**L**); leaf trichomes (**M**). Scale bar 200 µm.

Detailed analysis of *pbZIP23::GUS* seedlings showed GUS expression in the root meristematic region and lateral root primordia (**Fig. 4A-D**). In the differentiation zone of the primary root, there was no visible GUS expression in the 7-day-old seedlings (**Fig 4B-D**), but it became detectable, although weak, in the vasculature of 12-day-old, seedlings (**Fig. 4I,J**), and was stronger in 14-day-old seedlings (**Fig. S3**). The root hairs remained without GUS staining in younger or older seedlings (**Fig. 4D,J**). In the hypocotyl, there was a weak GUS signal in the vascular cylinder (**Fig. 4E,F**), whereas in the cotyledons there was a clear signal in the vascular tissues (**Fig. 4G**) that was present in the mesophyll cells in older seedlings as well (**Fig. 4L, S3**). The shoot apical meristem and leaf primordia showed a strong GUS expression (**Fig. 4E,H**), which was also observed in the vascular tissue and mesophyll of leaves from older seedlings (**Fig. 4M, S3**). The leaf trichomes, however, did not have GUS expression (**Fig. 4M**). This analysis showed that *bZIP23* is expressed in the root and shoot apical meristems, and it is not expressed in root hairs and leaf trichomes, showing also some expression in root vasculature and mature leaves.

**Figure 4.**
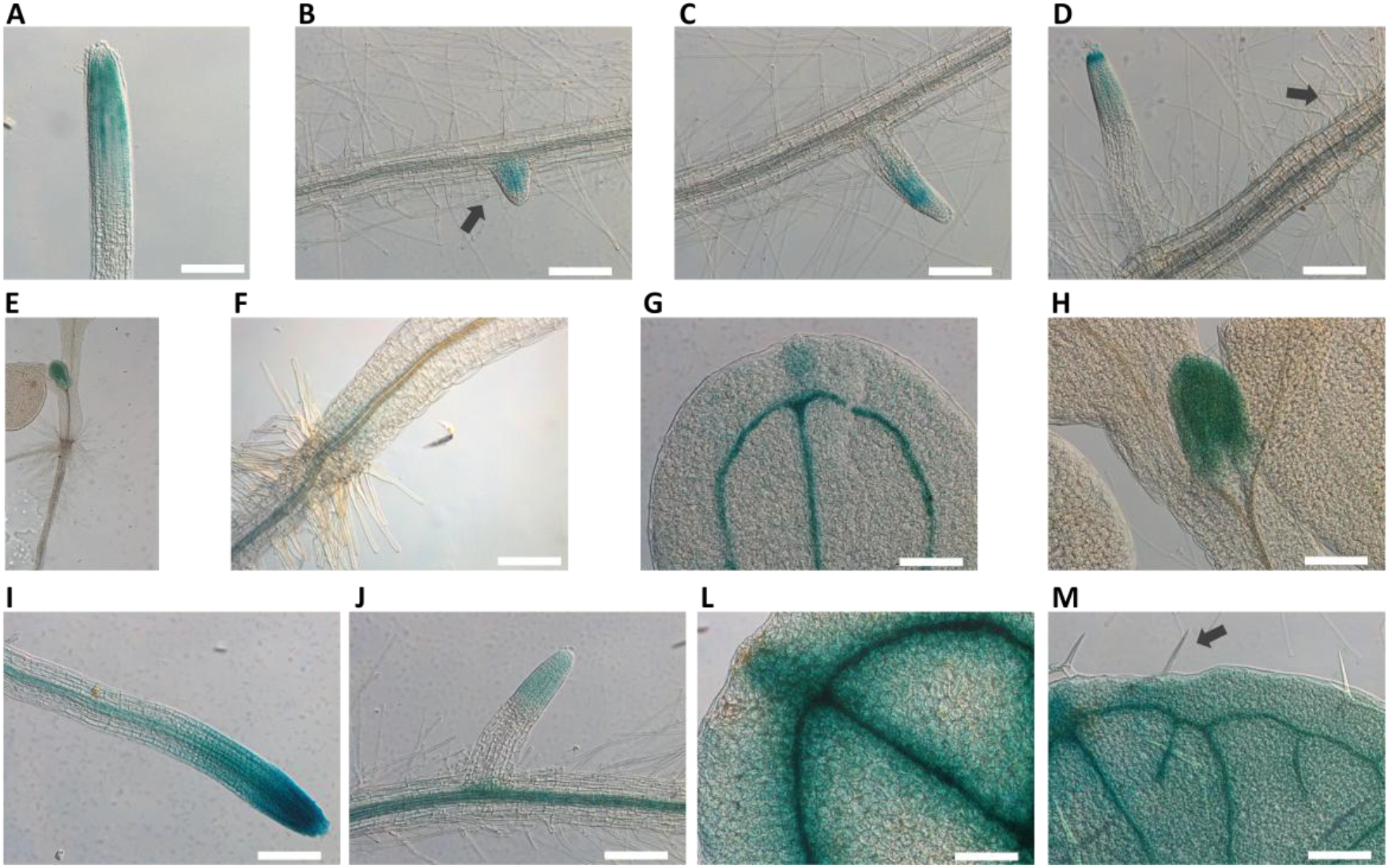
Analysis of *pbZIP23::GUS* seedlings after histochemical GUS staining. Seven-day-old seedlings grown on MS (control) showing the root tip (**A**); lateral root primordia (**B**,**C**); root maturation zone with root hairs and lateral root (**D**); root and base of hypocotyl (**E**,**F**); cotyledon (**G**); shoot apical meristem and leaf primordia (**H**); 12-day-old seedlings showing root apex and elongation zone (**I**); lateral root (**J**); cotyledon (**L**); leaf trichomes (**M**). Scale bar 200 µm.

Overall these results indicate that *bZIP19* and *bZIP23* have different, and mostly non-overlapping, tissue-specific expression patterns (**Fig. 3**,**4**). GUS expression indicates that only *bZIP23* is expressed in meristematic tissue, and is less expressed elsewhere, while *bZIP19* is expressed throughout the seedling in most differentiated cells and tissues.

### Tissue-specific expression patterns of *bZIP19* and *bZIP23* in reproductive organs

To further investigate the expression patterns of *bZIP19* and *bZIP23* in Arabidopsis, we analysed flowers and siliques of soil-grown *pbZIP19::GUS* and *pbZIP23::GUS* lines. Analysis of *pbZIP19::GUS* flowers showed GUS expression at the base of the flower, in the receptacle next to the peduncle (**Fig. 5A**), which remained expressed in this region in the peduncle-silique node at the later stages of silique development (**Fig. 5B-D**). It also showed GUS staining in the filament of the stamen and in the vascular tissues of the petals (**Fig. 5A**). In the silique, there was GUS expression in the septum and in the funiculus, which connects the seed to the placenta (**Fig. 5C,D**). On the other hand, analysis of *pbZIP23::GUS* flowers showed GUS expression in the stamen filament and anthers, and no expression in the flower receptacle (**Fig. 5E,F**). The peduncle-silique node had a very weak GUS signal (**Fig. 5E-G**), and no signal was detected in the silique structures (**Fig. 5G,H**). These results indicate a different expression pattern between *bZIP19* and *bZIP23* in flowers and siliques, with *bZIP19* expressed in different structures of flowers and siliques whereras *bZIP23* is mainly expressed in the stamen.

**Figure 5.**
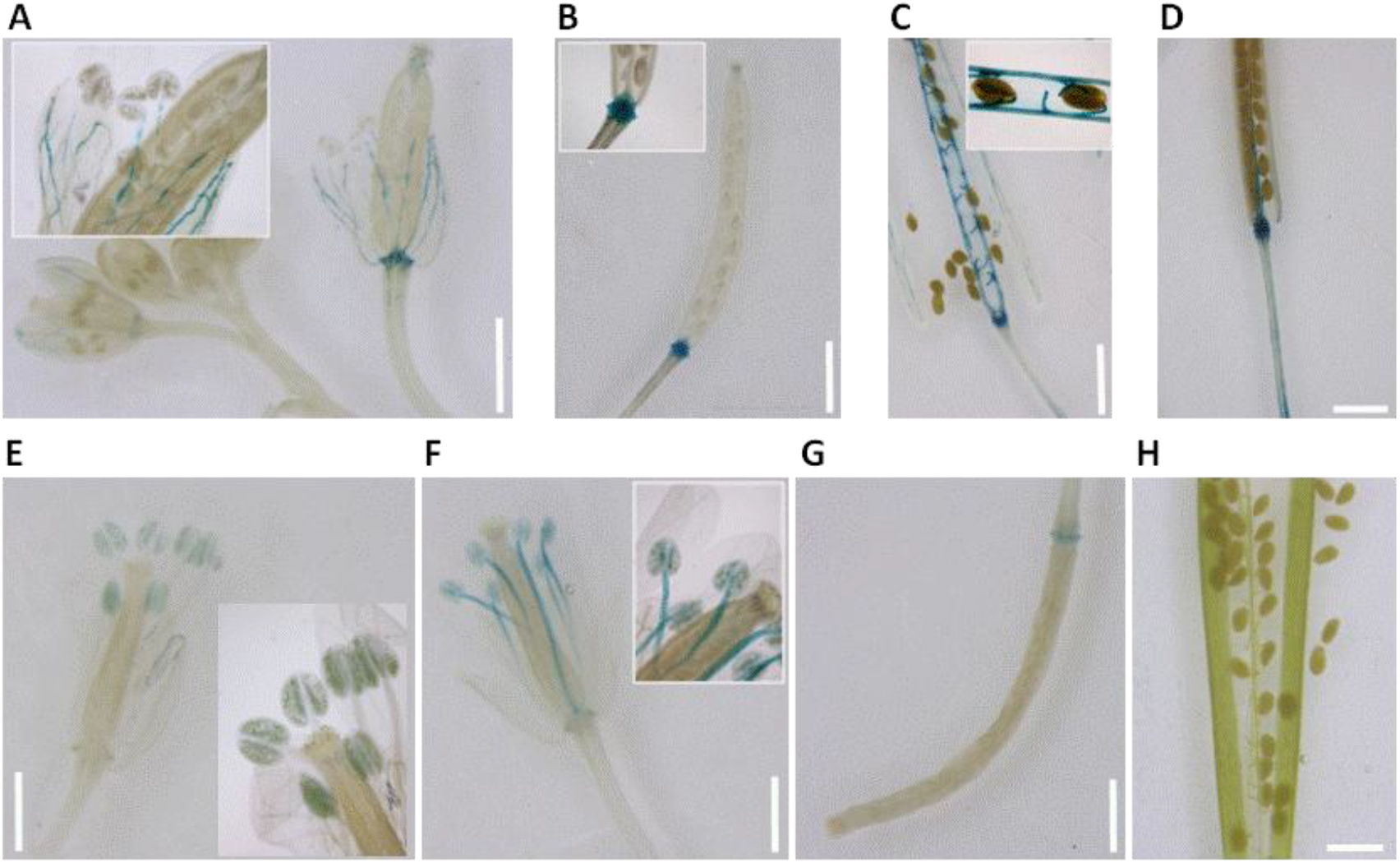
Histochemical GUS analysis of Arabidopsis flowers and siliques. Soil-grown *pbZIP19::GUS* plants (**A-D**) showing flowers (**A**), young silique (**B**) and mature silique (**C**,**D**) and soil-grown *pbZIP23::GUS* plants (**E-H**) showing flowers (**E**,**F**), young silique (**G**) and mature silique (**H**). Scale bar = 1 mm.

## DISCUSSION

We used *bzip19* and *bzip23* single and double mutants and *pbZIP19::GUS* and *pbIP23::GUS* Arabidopsis lines to investigate the individual contribution of bZIP19 and bZIP23 transcription factors as regulators of the Zn deficiency response. Comparison of rosette phenotype, dry weight and Zn content between mutants and wild-type plants grown under Zn deficiency showed the absence of phenotype in *bzip23*, suggesting compensation by its homologue bZIP19, and showed a strong Zn deficiency phenotype in *bzip19*, suggesting that bZIP19 is not compensated by bZIP23. In addition, the Zn deficiency hypersensitive phenotype of *bzip19/23* was an enhanced phenotype compared to that of *bzip19* showing that both transcription factors, bZIP19 and bZIP23, are necessary for the regulation of the Zn deficiency response (**Fig. 1**). Therefore, although bZIP19 and bZIP23 were shown to have functional redundancy (Lilay *et al*. 2019), our analysis of single and double mutant phenotypes showed that, using the nomenclature proposed by Briggs *et al*. (2006), bZIP19 and bZIP23 are unequally redundant.

To obtain further insight on the individual contributions of bZIP19 and bZIP23, we analysed tissue-specific gene expression patterns of *bZIP19* and *bZIP23* using *pbZIP19::GUS* and *pbZIP23::GUS* Arabidopsis lines. Previous work with these lines (Lilay *et al*. 2019) showed that *bZIP19* is mostly expressed in roots while *bZP23* seems to be more expressed in leaves. Here, a more detailed analysis revealed that *bZIP19* and *bZIP23* have different, and mostly non-overlapping, tissue-specific expression patterns (**Fig. 2-4**). GUS histochemical analysis indicated that *bZIP19* is expressed along the mature root. It appeared expressed across the epidermis, cortex and vascular cylinder, although thorough root cross-section observations are needed for confirmation. GUS staining also showed *bZIP19* expression in root hairs along the mature root. These are specialized elongated cells that extend the radial reach of roots and originate from differentiated epidermis cells (Ramakrishna & Barberon 2019). GUS staining analysis of the root also indicated that, on the other hand, *bZIP23* is mostly expressed in the apical meristem of primary and lateral roots, and is only expressed in root vascular tissue in older seedlings (12- and 14-day-old *pbZIP23::GUS* seedlings but not 7-day-old), suggesting that developmental cues play a role in regulating *bZIP23* expression. These different expression patterns of *bZIP19* and *bZIP23* in the root indicate that the bZIP19 transcription factor is responsible for sensing cellular Zn status and regulating the Zn deficiency response in the mature root and root hairs, where most nutrient acquisition and uptake takes place (Marschner 2012). This may explain, at least partly, the absence of phenotype in *bzip23* and a strong Zn deficiency phenotype in *bzip19* (**Fig. 1**).

GUS histochemical analysis in *pbZIP19::GUS* also indicated that *bZIP19* is expressed in the root-hypocotyl node, at the peduncle-flower receptacle node and peduncle-silique node (**Fig. 3**), that is, at transport junctions between organs. *bZIP19* expression was also observed in the silique (**Fig. 5**), in the region of the septum and replum, in between the two valves, and in the funiculus, which is the stalk that attaches the seed to the placenta (Herrera-Ubaldo & de Folter 2022). Thus, *bZIP19* is consistently expressed in tissues and organs involved in nutrient uptake and transport pathways, from the root throughout the plant and delivery to the seed. This highlights that Zn homeostasis regulation and Zn sensing are important in these structures, and indicates that such regulation relies mainly on the bZIP19 transcription factor. This may additionally explain the strong phenotype of *bzip19* mutant and its reduced shoot/root Zn concentration ratio at Zn deficiency (**Fig. 1**). We previously observed that soil-grown *bzip19/23* double mutant, although not showing developmental penalties compared with the wild-type, had significantly lower seed Zn concentration, possibly reflecting its limitations in coping with fluctuating Zn availability in soil (Huizinga *et al*. 2024). The observed reduced seed Zn content in *bzip19/23* would be in line with a role of bZIP19 in seed Zn filling.

GUS analysis in *pbZIP23::GUS* lines also indicated *bZIP23* expression in leaf vascular tissues and mesophyll, but, similarly to observations in the root, only detected in seedlings at older developmental stages (12- and 14-day-old). In the leaf trichomes, similarly to the root hairs, there was only *bZIP19* and not *bZIP23* expression (**Fig. 3**,**4**). In Arabidopsis, trichomes are large non-glandular epidermal cells with a characteristic star-shaped architecture, and can be found in most aerial organs (Ishida *et al*. 2008). Both trichomes and root hair cells, which showed GUS staining in *pbIP19::GUS* seedlings, result from cell patterning processes occurring in the epidermis (Ishida *et al*. 2008). In Arabidopsis, there is Zn accumulation at the base of trichomes which correlates with Zn concentration in leaves, suggesting that trichomes play a role in Zn sequestration and detoxification (Ricachenevsky *et al*. 2021). Likewise, accumulation of Zn in trichomes was also shown in the metal hyperaccumulator and hypertolerant *Arabidopsis halleri* (Sarret *et al*. 2009). The expression of *bZIP19* in trichomes indicates that Zn sensing and homeostasis regulation is important in these cells, suggesting that trichomes may have a more dynamic role than merely of a storage cell with regard to Zn homeostasis.

Comparing the *bZIP19* and *bZIP23* expression patterns inferred from the analysis of *pbZIP19::GUS* and *pbIP23::GUS* lines, it can be concluded that *bZIP19* is generally more expressed than *bZIP23*, and that while *bZIP19* is expressed throughout the plant in most differentiated tissues, *bZIP23* is mainly expressed in the root and shoot apical meristems (**Fig. 4**). The observed GUS staining in meristematic cells highlights the importance of Zn homeostasis regulation and adequate Zn availability in these cell types, and indicates that, rather than bZIP19, it is bZIP23 transcription factor that is responsible for Zn sensing and regulation in meristems. The activity of meristems is controlled by auxin and cytokinin dynamics (Salvi *et al*. 2020), which suggests that phytohormone signaling may be involved in regulating bZIP23 activity. Future work should investigate how phytohormones influence bZIP23 activity, and how bZIP23 activity affects meristematic function.

bZIP19 and bZIP23 have a high degree of amino acid sequence identity, cluster together phylogenetically and are likely two paralogs originating from a previous whole genome duplication event in the Brassicaceae (Castro *et al*. 2017). Whole genome duplications and small-scale duplications are key events for evolution. Whole genome duplications have been abundant in plant evolution, and are thought to contribute to increased plant diversity and adaptation (Almeida-Silva & Van de Peer 2023). The fate of paralogous gene pairs derived from duplication events may comprise one copy undergoing nonfunctionalization, if it has no selective pressure, or retention of gene copies. If the retained duplicate gains a new function there is neofunctionalization, or if the function of the ancestral gene is distributed between the two copies there is subfunctionalization (Defoort *et al*. 2019). Our results suggest subfunctionalization of *bZIP19* and *bZIP23*, where their differential tissue-specific expression pattern explains their evolutionary retention.

Additional details of the individual function of bZIP19 and bZIP23 paralogs, which would further clarify their evolutionary retention, have yet to be understood. The target gene set identified with the analysis of *bzip19/23* mutant includes several *ZIP* transporter family members (*ZIP1,3,4,5,9,10,12,IRT3*) (Assunção *et al*. 2010), but gene expression analysis in *bzip19* and *bzip23* single mutants suggested that *ZIP9* target gene expression is mainly under control of bZIP19 whereas *ZIP12* target gene is under control of bZIP23 (Inaba *et al*. 2015). It could be suggested that the regulation of Zn homeostasis by bZIP23 in meristematic cells would require a different repertoire of *ZIP* transporter target genes compared with the regulation of Zn homeostasis in differentiated cells by bZIP19. In addition to differences in the repertoire of target gene sets, differences between *bZIP19* and *bZIP23* promoter cis-regulatory elements, and eventually tissue-specific protein partners may further clarify the individual function of bZIP19 and bZIP23 transcription factors.

## CONCLUSION

bZIP19 and bZIP23 are key regulators of the Zn deficiency response in Arabidopsis and act as sensors of cellular Zn status. They are likely paralogs and we are just starting to understand their unequal functional redundancy. This study advances knowledge on the individual contribution of each transcription factor through the analysis of *bZIP19* and *bZIP23* gene expression patterns using *pbZIP19::GUS* and *pbZIP23::GUS* lines. *bZIP19* is generally more expressed than *bZIP23*. It is expressed in the root and throughout the plant, in differentiated cells and tissues, including those involved in transport pathways, whereas *bZIP23* is mostly expressed in meristematic tissue. These different tissue-specific gene expression patterns may explain the unequal redundancy between bZIP19 and bZIP23, and also, their evolutionary retention possibly through subfunctionalization. This knowledge has implications for the mechanistic understanding of the Zn sensing and deficiency response in Arabidopsis and also highlights the individual contributions of F-bZIP transcription factors in the pursuit of improved Zn nutritional value and use-efficiency in crops.

## AUTHOR CONTRIBUTIONS

A.C., G.H.L., M.B., P.H.C., and A.G.L.A. designed and performed experiments, and analysed data. A.C. and A.G.L.A. wrote the manuscript, revised by all authors.

## Supporting Information

**Figure S1.**
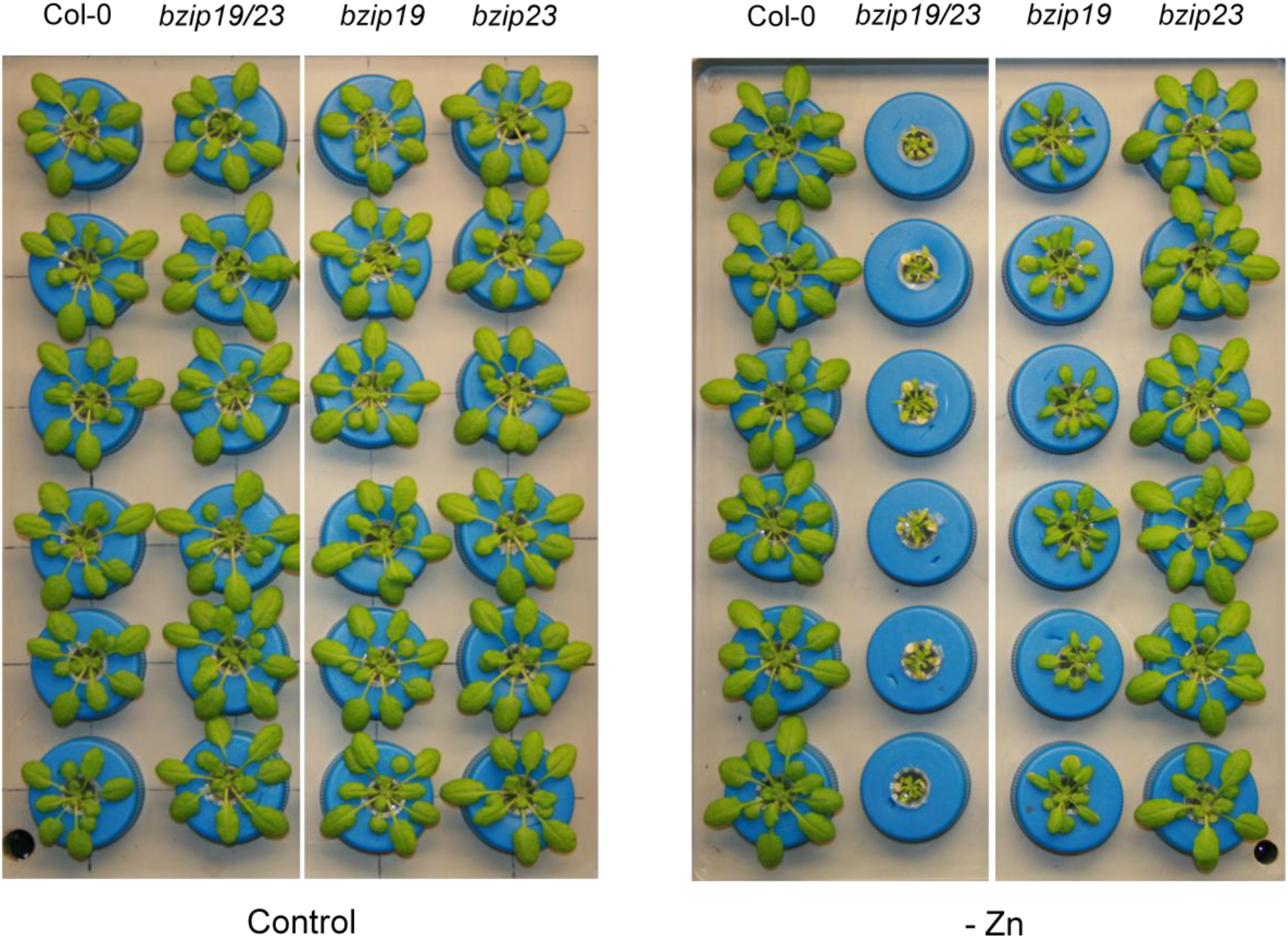
Phenotype of Arabidopsis *bzip19, bzip23, bzip19/23* and Col-0 plants grown in hydroponics with control or Zn deficient (-Zn) nutrient solution. Image shows rosettes (6 plants per genotype) from 4-week-old plants.

**Figure S2.**
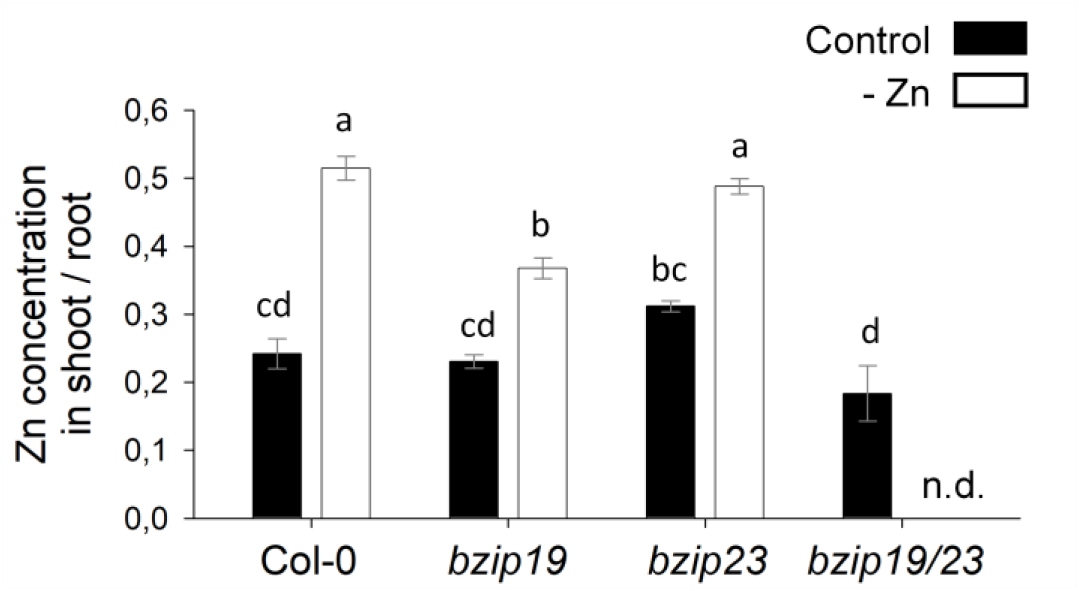
Ratio between shoot and root Zn concentration in *bzip19, bzip23, bzip19/23* and Col-0 Arabidopsis plants grown in hydroponics with control or Zn deficient (Zn-) nutrient solution. Shoot/root ratio was calculated from Zn concentration (µg g^-1^ d.w.) measured by ICP-OES, mean ± s.e., n= 3 to 5. There is no data (n.d.) for *bzip19/23* roots. Different letters indicate significant differences (P < 0.05), determined using one-way analysis of variance followed by Tukey post hoc test.

**Figure S3.**
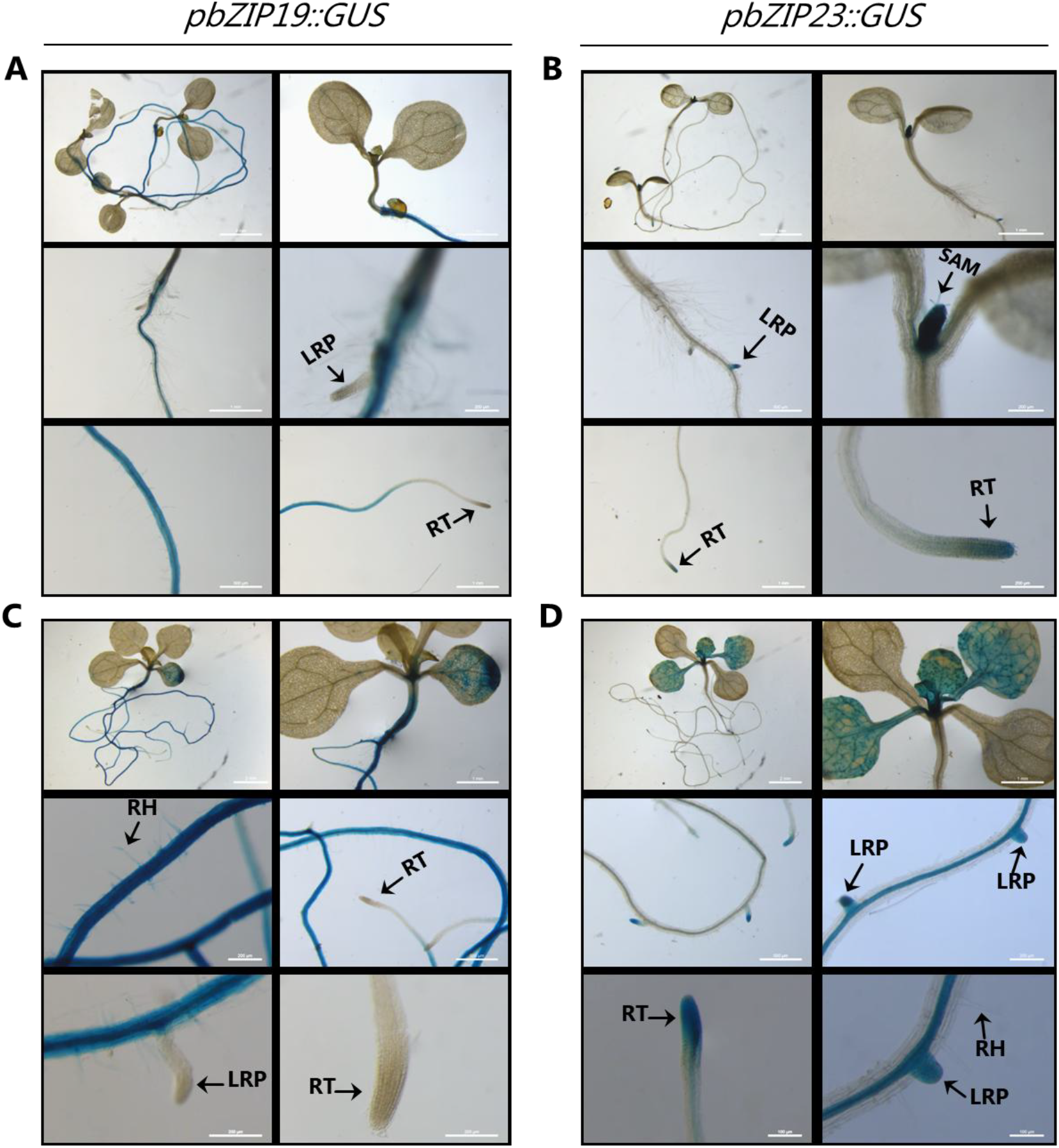
Analysis of *pbZIP19::GUS* and *pbZIP23::GUS* seedlings after histochemical GUS staining. (**A**,**B**) Seven-day-old seedlings grown on MS (Zn-); (**C**,**D**) 14-day-old seedlings grown on MS (Zn-). Arrows are pointing at RT (root tip), SAM (shoot apical meristem), LRP (lateral root primordia) and RH (root hairs).

